# Genetic Variability Analysis of Hot Pepper(Capsicum annuum.L) at Haramaya University, Eastern Ethiopia

**DOI:** 10.1101/2024.11.07.622433

**Authors:** Abdurazak Sufiyan

## Abstract

Hot pepper is a globally important spice utilized for food flavoring, enhancement, and coloring, with a growing demand over time. In the 2022 cropping season, this study aimed to assess the genetic variability among 19 landrace genotypes and one improved hot pepper genotype under semi-irrigation conditions. Twelve quantitative parameters were evaluated using a randomized complete block design with three replications. Analysis of variance revealed highly significant differences (P ≤ 0.01) for all traits except days to harvest and fruit weight, which were non-significant. Coefficients of variation ranged from 2.92 for days to fifty percent flowering to 31.47 for number of fruits per plant, indicating substantial variability among accessions. Fruit length varied from 4.5cm (accession 229694) to 11.17cm (accession 229699), while fruit width ranged from 8.33cm to 31.4cm, and fruit weight from 11.25g to 24.88g. Accession 229697 exhibited the highest fruit number per plant (42.67), while M/fana showed the lowest (12.67), with a mean value of 23.64. Fruit yield per plot ranged from 0.26kg/plot (accession 229696) to 0.91kg/plot (accession 28337), with approximately 40% of genotypes surpassing the population mean (0.5935kg/plot). High phenotypic and genotypic coefficients of variation, along with high genetic advance and heritability, were observed for yield per plot, number of fruits per plant, and fruit width, suggesting potential for improvement through selection. Based on the study results, accession 28337 for yield per plot, accession 229697 for fruit number per plant, and accession 229695 for fruit width are recommended for future selection.

## Introduction

According to [1], hot pepper ranks as the second most important vegetable worldwide, just behind tomatoes. It is the most widely used spice, adding flavor, color, and essential nutrients to food. Grown extensively throughout the tropics, hot pepper is an important spice crop. Although over 100 species fall under the genus Capsicum, most experts focus on just two primary species: *Capsicum annuum L*. and *Capsicum frutescens L*. [2]. The genus, commonly referred to as red chile, hot red pepper, chili pepper, tabasco, paprika, and cayenne, belongs to the Solanaceae (nightshade) family.

Distinguishing hot pepper from closely related species, such as *C. chinense* (noted for its intense heat) and *C. frutescens* (tabasco pepper), can be challenging due to overlapping morphological characteristics. These three species share a common ancestral gene pool, leading to confusion in naming, with terms like pepper, chili, chile, aji, paprika, and capsicum often used interchangeably [3].

Hot pepper holds significant nutritional value, particularly as a source of vitamins A and C, and provides a substantial portion of Ethiopians’ daily vitamin intake, with an average per-person consumption of around 15 grams daily. In Ethiopia, hot pepper is cultivated on roughly 246,000 hectares, mostly on small farms, yielding about 400 kg/ha of dry fruit on average [4].

Despite its importance, hot pepper production remains low, with average yields of 7.6 t/ha for green pods and 1.6 t/ha for dry pods [5, 6, 7]. This low productivity is due to suboptimal varieties, poor farming practices, and widespread fungal, bacterial, and viral diseases [6]. According to [8], rising demand for chili pepper in fresh and industrial markets has increased the need for high-yield, visually appealing, and commercially viable varieties. Developing high-yielding chili varieties with desirable traits could improve overall productivity.

Genetic diversity within the crop is key to its improvement, with genetic variability serving as the foundation for enhancing crop characteristics [9]. In Ethiopia, several studies have examined hot pepper’s genetic diversity using morphological markers for academic and research purposes [10-16]. One such study at Dire Dawa University’s Tony Farm Site [17] found that current accessions had not been characterized under highland conditions, which differ significantly from Dire Dawa’s lowland environment. This study therefore aims to morphologically characterize hot pepper accessions for these distinct conditions.

## Methodology

### Description of study area

The study area of these field experiments was found in Haramaya Woreda as shown in the map below Haramaya (Fig 1). Haramaya University is located in the Haro Maya district, East Hararghe Zone of the Oromia Regional State in Ethiopia. Haramaya University is situated approximately 510 kilometers (320 mi) east of Addis Ababa, the capital city of Ethiopia.

**Fig 1.**
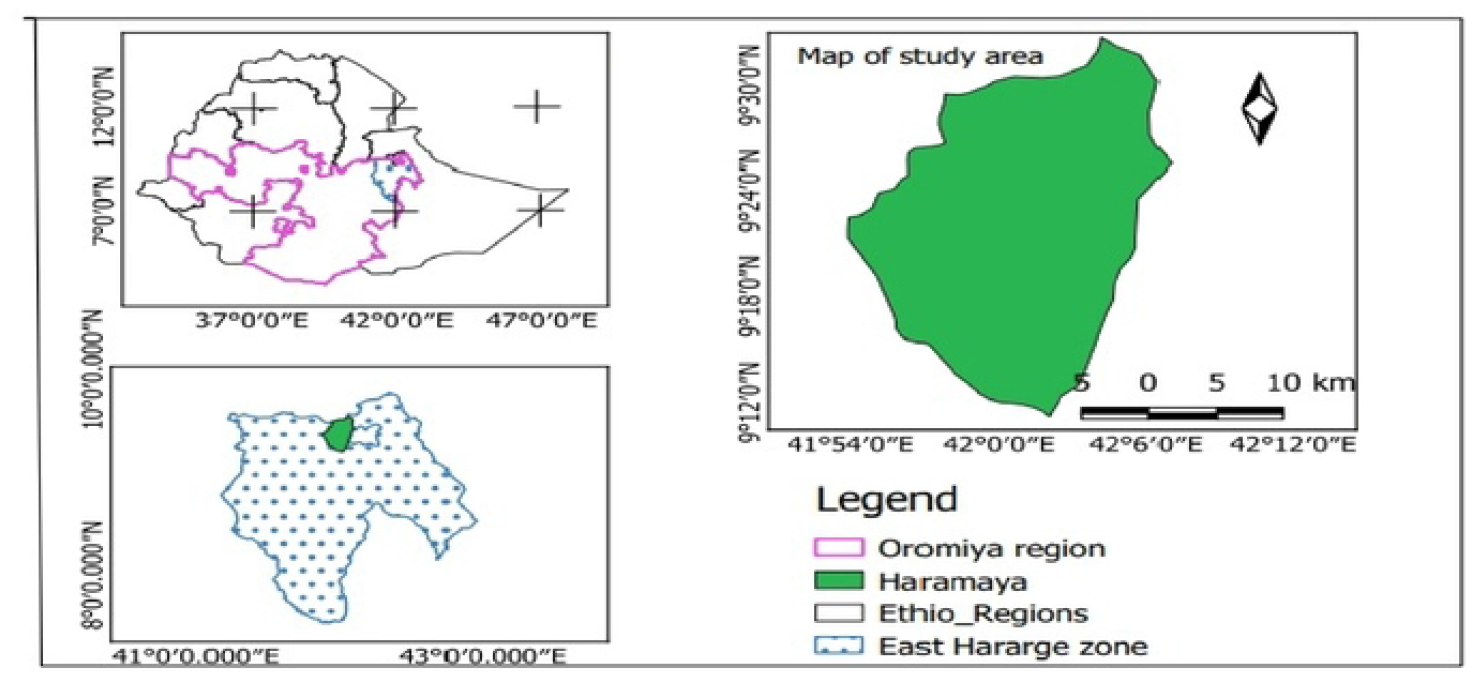
Map of study area.

The rare research site is located at 9 o26’ N latitude, 42 o3’ E longitudes at an altitude of 1980 m.a.s.l. The mean annual rainfall is 760 mm [18]. The mean annual temperature is 16 °C. The mean relative humidity is 50%, varying from 20 to 81%. The soil of the experimental site is alluvial type with an organic carbon content of 1.15%, total Nitrogen content of 0.11%, available Phosphorus content of 18.2 mg kg soil-1, the exchangeable Potassium content of 0.65 cmolc kg soil-1, pH of 8.0 and percent sand, silt, and clay content of 62.92, 19.64, and 17.44, respectively [19].

Planting materials used for this study shown in Table 1 below comprised 20 genotypes among which 19 were treatment tests with landrace genotypes maintained at EBI (Ethiopia Biodiversity Institute) and the other genotypes was Marako fana (improved variety) used as check varieties which were obtained from Fedis Agricultural Research Center.

**Table 1.** Passport of experimental material used for this study was obtained from EBI.

| Accession No | Region/Region | Zone/Zone | Woreda | Latitude | Longitude | Altitude |
| --- | --- | --- | --- | --- | --- | --- |
| 20206 | Oromiya | Misrak Wellega | Sasiga | 09-12-47-N | 36-25-52-E | 1667 |
| 20207 | Oromiya | Misrak Wellega | Wama hagelo | 08-48-31-N | 36-55-12-E | 1597 |
| 20212 | Oromiya | Illubabor | Metu | 08-20-07-N | 35-35-08-E | 1678 |
| 20213 | Oromiya | Illubabor | Ale | 08-11-36-N | 35-32-27-E | 1758 |
| 20214 | Oromiya | Illubabor | Chora | 08-21-53-N | 36-03-13-E | 1652 |
| 27845 | South Ethiopia Region | Bench maji | Gurra farda | 06-46-37-N | 35-11-50-E | 1408 |
| 27846 | South Ethiopia Region | Bench maji | Gurra farad | 06-90-46-N | 35-18-32-E | 1121 |
| 28334 | Oromiya | Illubabor | Harrumu | 08-21-36-N | 35-43-39-E | 1657 |
| 28336 | Oromiya | Illubabor | Durame | 08-26-45-N | 35-52-54-E | 1868 |
| 28337 | Oromiya | Illubabor | Durame | 08-25-43-N | 35-52-48-E | 1877 |
| 28338 | Oromiya | Illubabor | Algae | 08-84-43-N | 35-45-60-E | 1831 |
| 229694 | Benishangul Gumuz | Metekel | Dibate | - | - | 1640 |
| 229695 | Benishangul Gumuz | Metekel | Dibate | - | - | 1650 |
| 229696 | Benishangul Gumuz | Metekel | Dibate |  |  | 1700 |
| 229697 | Benishangul Gumuz | Metekel | Dibate | - | - | 1440 |
| 229698 | Benishangul Gumuz | Metekel | Dibate | - | - | 1520 |
| 229699 | Amhara | Misrak gojam | Bibugn | 11-07-00-N | 37-44-00-E | 1850 |
| 229700 | Amhara | Misrak gojam | Bibugn | 11-06-00-N | 37-44-00-E | 1830 |
| 229701 | Amhara | Misrak gojam | Hulet ej<br>enese | 11-05-00-N | 37-46-00-E | 1940 |
| Melka Fana |  |  |  |  |  |  |

The experimental design was a randomized complete block design (RCBD) with three replications. Seeds of each genotype were sown in April 2022 on a seedbed size of 7.8m×2m with a total seed bed area of 15.6m^2^ (each genotype was sown on two rows of 2m long). Transplanting to the actual field was done when the seedlings attained 20 to 25 cm height and or 40 days after sowing. Each seedling genotype was planted on a plot size of 1.5 m x 2m (with a total plot size of 3 m^2^) and the distance between plots and between replication was 0.7m and 1m respectively. Each plot within a replication consists of four rows and each row contains fivplants with a total of 20 plants per plot. The Seedlings were spaced 50 cm between plants and 70 cm between rows. The experimental plots will be fertilized with 200 kg/ha DAP as a side dressing during the transplanting operation in addition, 100 kg/ha UREA, half of it during the transplanting and half of it 15 days after transplanting will be applied [20].

### Experimental Procedure

#### Land preparation

Larger clods were broken into small particles and finally attained into a desirable tilth to ensure proper growing conditions. Recommended doses of well-decomposed cow dung, manure, and chemical fertilizers were applied and mixed well with the soil of each plot. Proper irrigation and drainage channels were also prepared around the plots. Each unit plot was prepared keeping 5cm height from the drains. The bed soil will be made friable and the surface of the bed be leveled.

#### Planting

In April 2022, seeds of 19 landrace genotypes and 1 check, totaling 20 genotypes, were sown on a seed bed. The bed was initially covered with dry grass for 20 days, followed by the application of raised shade to shield the seedlings from intense sunlight until they were ready for transplanting. After 55 days of seeding, robust and healthy seedlings were carefully selected and transplanted into the well-prepared field. Each hole received one seedling, and after planting, the bases of the seedlings were covered with soil and firmly pressed by hand. Four days before planting capsicum seedlings, a mixture of well-decomposed cow dung, TSP, and other fertilizers was applied to the plots and thoroughly incorporated into the bed soil. During the final bed preparation, one-fourth of both Urea and MP fertilizers were applied. The remaining Urea and MP were top-dressed in three equal installments, 30, 45, and 60 days after planting.

#### Irrigation

during the initial phase of planting seed on the seed bed, irrigation was used. After that semi-irrigation was used based on the availability of rain.

#### Cultural practices

mulching, weeding, cultivation, watering, and earthing-up were done at the appropriate time to facilitate root, to control disease infestation, and to control waterlogging. Integrated Weeding and hoeing were done to improve soil structure and reduce competition of weeds and earthing-up was done as required to prevent exposure of roots to direct sunlight.

#### Harvesting

Harvesting of fruits was started at 75 DAP and continued up to 25 DAP with an interval of 25 days. Harvesting was done usually by hand. Five plants from each row or plot left the plants growing at both ends of each row to avoid edge effects, were harvested to estimate fruit yield and other yield-related parameters.

### Data collection

Quantitative (12) morphological data was collected according to the descriptor for Capsicum [21]. At harvest, 10 guarded plants were randomly taken from each plot to measure quantitative morphological character. Some of the characters were measured before harvest. The sampling was done in such a way that the border effects were completely avoided. For this purpose, the outer two lines and the end of the middle rows were excluded.

The following quantitative morphological data was collected: **Plant height (PH):** Length in centimeters of the central axis of the stem, measured from the soil surface up to the tip of the stem, and the average was recorded. Recorded when in 50% of the plants the first fruit has begun to ripen. **Days to 50% flowering (DFL):** Number of days from transplanting to when 50% of plants in a plot open the flower. **Days to 50% Fruiting (DF):** Number of days from transplanting until 50% of the plants bear mature fruits at the first and second bifurcation. Recorded on mature fruits. **Number of flowers per axil (NFLA):** the number of flowers counted per axil recorded on fully open flower. **Days to first harvest (DH):** Number of days from transplanting to first harvest. **Number of fruits per plant (NFP):** Average number of chili fruits, counted at harvest on 10 sample plants of each plot. **Fruit length (FL):** The average length of five chili fruits was measured in centimeters on 10 plants of each plot. **Fruit width (FW):** Measured at the widest point. Average fruit width of 10 ripe fruits. **Fruit Weight (FWT):** Average fruit weight of 10 ripe fruits of the second harvest. **Number of seeds per Fruit (NSF):** Average of at least 10 fruits selected from 10 random plants. **1000-seed weight [g] (TSW):** The weight of 1000 seeds is in measured each plot. **Yield per plot [Kg] (FYPP):** The weights of total fruits harvested in each plot from all central row plants were recorded to estimate yield per plot.

### Data Analysis

#### Descriptive Statistics

The mean value of each character understudy was summarized using Microsoft Excel and subjected to analysis of variance following the procedure described by [22] and [23].

#### Estimation of genetic parameter

##### Genotypic and phenotypic coefficient of variation

The phenotypic and genotypic variability of each quantitative trait was estimated as phenotypic and genotypic variances and coefficients of variation. Phenotypic and genotypic components of variance were estimated by using the formula given by [24, 25]. Genotypic and phenotypic coefficient of variation was computed according to [26, 27, 25]

Genotypic variance (σ^2^g)

Where, σ^2^g = genotypic variance

Mg= mean square of genotype

Me = mean square of error

r = number of replications

Phenotypic Variance (σ^2^p) = σ^2^g + σ^2^e

Where, σ^2^g = Genotypic variance

σ^2^e = Environmental variance

σ^2^p = phenotypic variance

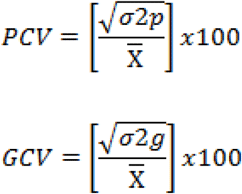

[28] classified the PCV and GCV estimates as follows:

Low, <10%

Moderate, 10-20%

High, >20%

##### Heritability

The broad sense heritability (H^2^) was estimated for all characters as the ratio of genotypic variance to the total or phenotypic variance as suggested by [25, 29, 30]

H^2^ = (σ^2^g/σ^2^p) x 100

Where, H^2^ = heritability in a broad sense

σ^2^p = phenotypic variance

σ^2^g = Genotypic variance

According to [31] heritability estimates in cultivated plants can be placed in the following categories.

Low, <30%

Moderate, 30-60%

High, >60%

##### Genetic Advance (GA) and Genetic Advance as a percent of the mean (GAM)

The genetic advance will be estimated according to [32, 33]:

GA = K *SDp* H^2^

Where, GA = Genetic advance

SDp = Phenotypic standard deviation on a mean basis;

H^2^ = Heritability in the broad sense.

k = the standardized selection differential at 5% selection intensity (K = 2.063).

GAM = [GA/PMC] x 100

Where GAM = Genetic advance as percent of mean

GA = Genetic advance

PMC = Populations mean character to be evaluated

The GA as a percent of the mean will be categorized as low, moderate and high as suggested by [32] as follows.

0 - 10% = Low

10 – 20 = Moderate

>20 = High

## Result and Discussion

### Range, mean, CV (%), and standard deviation of 20 genotypes

The mean performance of 20 hot pepper genotypes for 12 quantitative traits was detailed in S1 Table. Table 2 presents the mean, range, coefficient of variation (CV), and standard deviation values for these traits. Notably, accession 20206 exhibited the tallest plant height (64.27), while accession 229694 displayed the shortest (42.4). Days to fifty percent flowering ranged from 55 (accession 28337) to 83.33 (accession 20207), with a mean of 68.72.

**Table 2.**
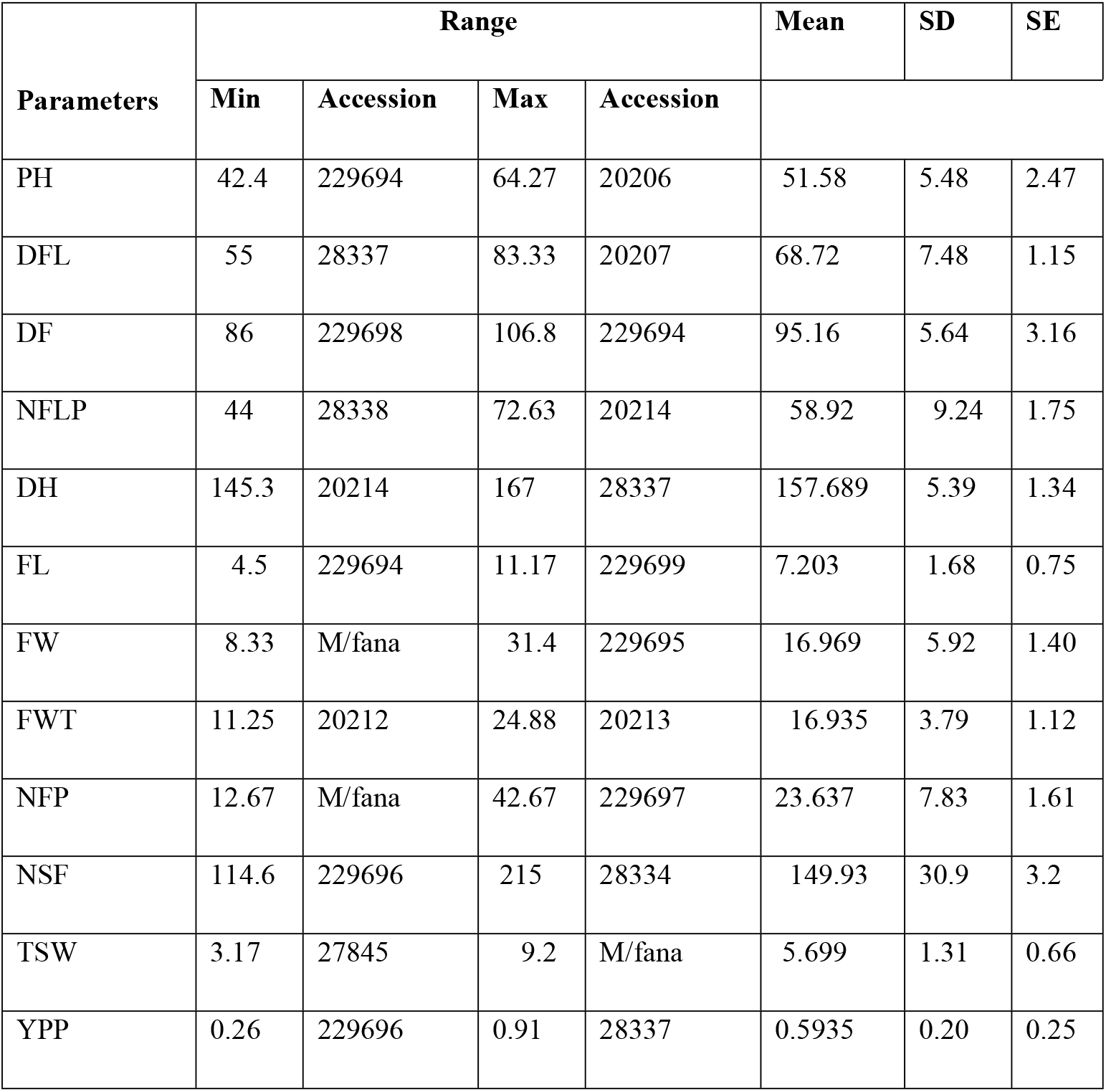
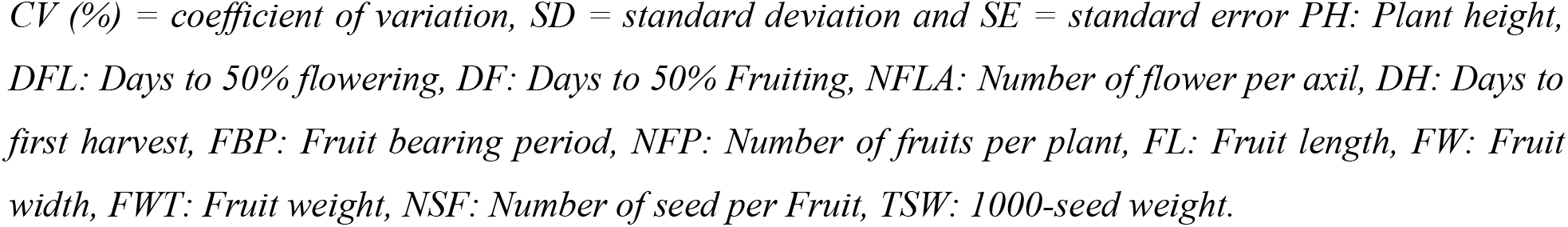
Range, mean, CV (%) and standard deviation of 20 genotypes.

Likewise, fruit-related traits showcased considerable variation, with accession 229694 recording the shortest fruit length (4.5cm) and accession 229699 registering the longest (11.17cm). Fruit width ranged from 8.33cm to 31.4cm, while fruit weight ranged from 11.25g to 24.88g. Accession 229697 boasted the highest fruit number per plant (42.67), contrasting with M/fana (12.67), which had the lowest. Fruit yield per plot ranged from 0.26kg/plot (accession 229696) to 0.91kg/plot (accession 28337), with a mean of 0.5935 kg/plot.

Approximately 40% of the genotypes outperformed the population mean. The findings align with similar studies conducted by [17, 16, 34] reflecting wide-ranging means for various traits and affirming the potential for pepper improvement.

### Analysis of variance (ANOVA)

Analysis of variance (ANOVA) was conducted for fruit yield per plot and other yield-related traits, revealing highly significant (P ≤ 0.01) mean squares for all traits except for days to harvest and fruit weight. The result of analysis of variance was presented in Table 3. The coefficient of variation was notably high for the number of fruit per plant (31.47) and lowest for days to fifty percent flowering (2.92), underscoring substantial variability among the accessions.

**Table 3.** Analysis of variance for different characters in hot pepper genotypes.

| Characters | Mean sum of square |  |  | CV |
| --- | --- | --- | --- | --- |
|  | Replication(r-1) = 2 | Genotype(g-1) = 19 | Error (r-1)(g-1) = 38 |  |
| Plant height | 3.870965 | 90.317731** | 18.273535 | 8.28 |
| Days to flowering | 0.772667 | 162.967579** | 3.983895 | 2.92 |
| Days to fruiting | 25.443167 | 95.409158** | 30.003342 | 5.75 |
| Number of flowers per plant | 0.686502 | 256.378658<sup>ns</sup> | 220.75880 | 25.21 |
| Days to harvest | 30.444435 | 87.358707<sup>ns</sup> | 52.892260 | 4.61 |
| Fruit length | 10.1051667 | 8.4525965** | 3.6770965 | 26.62 |
| Fruit width | 16.219852 | 105.109612** | 25.990246 | 30.04 |
| Fruit weight | 1.8435050 | 43.4944404<sup>ns</sup> | 27.084161 | 30.78 |
| Number of fruit per plant | 77.516667 | 175.483333** | 56.499123 | 31.47 |
| Number of seed per fruit | 1163.26667 | 2854.51228** | 380.61754 | 13.01 |
| Thousand seed weight | 8.92381500 | 5.19725754** | 2.2551746 | 26.35 |
| Yield per plot | 2.37049833 | 0.12476307** | 0.02261254 | 25.33 |

\*\* Denotes significant at 1% level of probability ns: non-significant

### Estimates of genetic parameter

#### Estimate of Genotypic and phenotypic coefficient of variation

Table 4 presents a comprehensive overview of variability concerning environmental, phenotypic, and genotypic factors, alongside heritability, genetic advancement, and coefficient variations. A clear trend emerges across traits: the phenotypic coefficient of variation (PCV) consistently exceeds the genotypic coefficient of variation (GCV), highlighting the strong influence of environmental factors. High PCV and GCV values, particularly for yield per plot, number of fruits per plant, and fruit width, point to significant potential for improvement through selective breeding. This difference between PCV and GCV across traits emphasizes the complex relationship between environmental factors and genetic variability. Furthermore, moderate to high heritability estimates add reliability to these findings.

**Table 4.** Estimate of genetic parameters.

| **Traits** | **PCV** | **GCV** | **h<sup>2</sup>(%)** | **GA** | **GAM(%)** |
| --- | --- | --- | --- | --- | --- |
| Plant height | 12.60 | 9.499 | 56.79 | 7.61 | 14.77 |
| Days to flowering | 10.98 | 10.59 | <u>93.01</u> | 14.48 | 21.08 |
| Days to fruiting | 7.56 | 4.91 | 42.08 | 6.24 | 6.57 |
| Number of flower per plant | 25.88 | 5.83 | **5.07** | 1.59 | 2.71 |
| Days to harvest | **5.08** | **2.14** | 17.83 | 2.95 | **1.87** |
| Fruit length | 31.84 | 17.50 | 30.22 | 1.43 | 19.86 |
| Fruit width | <u>42.64</u> | 30.26 | 50.36 | 7.51 | 44.30 |
| Fruit weight | 33.69 | 13.81 | 16.80 | 1.98 | 11.68 |
| Number of fruit per plant | 41.8 | 26.64 | 41.24 | 8.34 | 35.30 |
| Number of seed per fruit | 23.15 | 19.15 | 68.42 | <u>49.00</u> | 32.68 |
| Thousand seed weight | 31.53 | 17.37 | 30.34 | 1.12 | 19.73 |
| Yield per plot | 39.87 | <u>31.07</u> | 60.71 | **0.29** | <u>49.94</u> |

*Heritability (H%), genetic advance (GA), genetic advance as percent mean (GAM), phenotypic and genotypic coefficients of variation. Underlined and bolded are the maximum and minimum value respectively.*

Studies by [17, 34, 35] support these observations, reinforcing the robustness of the identified trends. The categorization of PCV and GCV into low, moderate, and high ranges by [28] further underscores these results. Notably, high PCV values are observed for several traits, including fruit width, yield per plot, fruit weight, and thousand seed weight. Moderate PCV values are seen for traits like plant height and days to flowering, while days to fruiting and days to harvest show low PCV values.

The high GCV values for yield per plot, fruit width, and number of fruits per plant, contrasted with moderate GCV for other traits, emphasize the potential for focused breeding. Similarly, studies on hot pepper by [17] confirm the findings on the number of fruits per plant and yield per plant, providing further evidence for the observed trends.

The noticeable differences between GCV and PCV underscore the significant role of environmental factors in trait expression. This discrepancy is particularly strong for traits such as fruit length, fruit weight, and thousand seed weight, reflecting a substantial environmental influence on these characteristics. Conversely, traits like days to flowering and days to fruiting show minimal differences between GCV and PCV, suggesting a reduced environmental impact.

Overall, the substantial genetic variability across traits indicates promising potential for targeted selection, supporting the enhancement of desirable traits and the development of improved cultivars.

#### Estimate of Heritability (%)

Heritability measures the proportion of phenotypic variance that is due to genetic factors, indicating the genetic component passed to future generations. Broad-sense heritability, which considers genotypic variance relative to total variance in a non-segregating population, provides insight into the potential effectiveness of selecting hybrids with specific traits [36, 37]. Traits with high broad-sense heritability suggest a substantial genetic contribution with limited environmental influence, making selection more effective. In contrast, traits with low heritability are harder to select for, as environmental factors play a larger role [38]. Heritability estimates, combined with genetic coefficients of variability as described by [27], offer valuable guidance on the expected gains from selection.

In this study, broad-sense heritability (H2) ranged from 5.07 for the number of flowers per plant to 93.01 for days to flowering (Table 4). Classifying heritability percentages into low, moderate, and high categories showed high heritability for days to flowering, number of seeds per fruit, and yield per plot. Meanwhile, previous studies [39, 40, 41, 42] report varying heritability for traits like branch number, fruit count per plant, fruit length, and primary branches per plant. High heritability for specific traits indicates that much of the phenotypic variation is due to genetic factors, allowing for reliable selection based on observable traits [43].

Moderate heritability was observed for traits including plant height, days to fruiting, fruit length, fruit width, number of fruits per plant, and thousand seed weight. Additionally, low broad-sense heritability estimates were recorded for traits such as the number of flowers per plant, days to harvest, and fruit weight, aligning with findings from Vijaya et al. [34].

#### Estimate of genetic advance and genetic advance % of the mean

Genetic advance measures the improvement achievable through selection within a population [44], making it a valuable indicator of the potential progress that can be expected from selection efforts [45].

The genetic advance as a percentage of the mean (GAM) at a 5% selection intensity is shown in Table 4, ranging from 1.87 for days to harvest to 49.94 for yield per plot.

According to [32], genetic advance as a percent of the mean can be categorized as follows: Low = 0-10%, Moderate = 10-20%, and High ≥ 20%. In this study, the highest GAM at a 5% selection intensity was observed for yield per plot, followed by fruit width, number of fruits per plant, days to flowering, and number of seeds per fruit. These findings are supported by [41], who reported similar results for dry yield per plant.

Low genetic advance was observed for days to harvest, number of flowers per plant, and days to fruiting, consistent with findings from [17] for days to fruiting. This is attributed to low phenotypic and genotypic coefficients of variation, underscoring the importance of genetic variability for selection-based improvement [34].

Heritable variation is more accurately assessed when heritability is evaluated alongside genetic advance. Since heritability alone does not indicate the expected progress from selection, it is most informative when paired with genetic advance estimates. Genetic advance is influenced by factors like selection intensity, heritability, and phenotypic variance. High genetic advance combined with high heritability suggests the presence of additive gene action [44]. In this study, high genetic advance paired with high heritability was noted for yield per plot, number of seeds per fruit, number of fruits per plant, fruit width, and days to flowering. This aligns with findings by [38] for the number of fruits per plant, and [34] for the number of fruits per plant and yield per plot. These traits show strong potential for selection, indicating that additive gene action is predominant and that direct phenotypic selection for these traits is effective.

## Conclusions

The current study revealed that yield per plot, number of fruits per plant, and fruit width exhibited high values of phenotypic coefficient of variation (PCV), genotypic coefficient of variation (GCV), and high heritability, along with significant genetic advance as a percentage of the mean. These findings suggest the predominance of additive gene action and a lesser influence of environmental factors on the expression of these traits, indicating the potential for improvement through selection. Furthermore, this implies the feasibility of enhancing these traits through selective breeding.

In conclusion, accession 28337 is recommended for yield per plot, accession 229697 for number of fruits per plant, and accession 229695 for fruit width based on the study’s results. Additionally, the following suggestions and recommendations are proposed:

1. The study observed high phenotypic and genotypic coefficients of variation, as well as significant genetic advance coupled with high heritability for yield per plot, number of fruits per plant, and fruit width, indicating the potential for improvement through selection. Emphasis should be placed on selecting genotypes with a higher number of fruits per plant, as this trait showed a significant positive correlation with yield per plot, suggesting its usefulness in selecting productive genotypes.
2. While morphological characterization was conducted in this study, it is recommended to consider molecular analysis for similar accessions to complement the findings. This integration of morphological and molecular studies could provide a comprehensive understanding of genetic variability.
3. The study was conducted with 19 accessions and 1 check variety. To broaden the scope of research, it is advised to include more accessions and varieties from diverse environments for further investigation. This would enhance the robustness and applicability of the study’s findings.

## Acknowledgement

The authors would like to express gratitude to the administrators of the Raare Research Station of Harmaya University and field data collectors for their cooperation.

## Supporting information

**S1 Fig. Map of study area**.

**S1 Table. Mean performance of 20 hot pepper genotype for 12 quantitative character**

## References

1. Rehima Mussema (2006). Analysis of Red pepper Marketing. The case of Alaba and Siltie in SNNPRS of Ethiopia. A Thesis submitted to Department of Agricultural Economics School of Graduate studies Haromany University. P: 1–8.

2. Purseglove, J.W. (1968). TropicaL crops, Dichotyledons. John Wiley and Sons, New York. Vol. 2: 524–530.

3. Zhang, Z., Lu, A. and D’arcy, W. G. Capsicum annuum Linnaeus, Special Plant, Flora of China,17, 313–313. 2002

4. CPI (Chile Pepper Institute) publication. New Mexico State University. 2007.

5. Alebachew Molla Nibret. Genetic Diversity of Hot Pepper (Capsicum annuum) from Selected Areas of Ethiopia Using Inter Simple Sequence Repeats (ISSR) Marker. A Thesis Submitted to Applied Biology Program in Partial Fulfillment of the Requirement of the degree of Master of Science in Biology (Biotechnology) School of Applied Natural Science Office of Graduate Studies Adama Science and Technology University. 2018.

6. Fekadu Moges. and Dandena, G. Status of Vegetable Crops in Ethiopia.Ugandan Journal of Agriculture; 2006. 12(2): 26–30

7. CSA. Area and Production of Crops (Private Peasant Holding, Meher Season). Addis Ababa. pp; 2006. 14–63.

8. Cabral N.S.S., Medeiros A.M., Neves L.G., Sudré C.P., Pimenta S., Coelho V.J., Serafim M.E. and Rodrigues R. Genotype x environment interaction on experimental hybrids of chili pepper. Genetics and Molecular Research 16 (2): 2017 gmr16029551. DOI 10.4238/gmr16029551

9. Pandit, M. K and Adhikary, S. Variability and Heritability Estimates in Some Reproductive Characters and Yield in Chilli (Capsicum annuum L.). International Journal of Plant and Soil Science; 2014 3 (7): 845–853.

10. Shumbulo A, Nigussie M, Alamerew S. Correlation and Path Coefficient Analysis of Hot Pepper (Capsicum annuum L.) Genotypes for Yield and its Components in Ethiopia. Adv Crop Sci Tech; 2017, 5: 277. doi: 10.4172/2329-8863.1000277

11. Belay Fasikaw and Tsehaye Yemane. Variability, Association and Path Coefficient Analysis of Green Pod Yield and Yield Components of Hot pepper (Capsicum annuum L.) Landraces at Mereb Lehke, Northern Ethiopia. Vol. 12(1), pp. 58–69, January-March 2020 DOI: 10.5897/JPBCS2019.0840 Article Number: 2C6A83763304 ISSN 2006-9758 http://www.academicjournals.org/JPBCS

12. Birhanu Habtie & Tiegist Dejene. Multivariate Analysis and Traits Association in Hot Pepper (Capsicum annuum) Landraces of Ethiopia. International Journal of Research Studies in Agricultural Sciences (IJRSAS), 2020; 6(10), pp. 42–51, 10.20431/2454-6224.0610005

13. Birhanu H, Tiegist D, Yigzaw D. Morphological Characterization of Hot Pepper (Capsicum annuum.L) Land Races of Ethiopia for Qualitative Characters”, International Journal of Research Studies in Science, Engineering and Technology, vol. 4, no. 9, pp. 4–9 2017.

14. Kahsay Y. Evaluation of Hot Pepper Varieties (capsicum species) for Growth, Dry pod Yield and Quality at M/Lehke District, Tigray, Ethiopia. International Journal of Engineering Development and Research. © 2017 IJEDR | Volume 5, Issue 3 | ISSN: 2321-9939

15. Belay F, Abate B, and Tsehaye Y. Genetic diversity studies for morphological traits of hot pepper (Capsicum annuum L.) genotypes in Central Zone of Tigray Region, Northern Ethiopia. Academic Journals Vol. 14(33), pp. 1674–1684, October, 2019. DOI: 10.5897/AJAR2019.14256. ISSN: 1991-637X. http://www.academicjournals.org/AJAR

16. Shimeles Aklilu, Bekele Abebie, Dagne Wogari and Adeferis TW. Genetic Variability and Association of Characters in Ethiopian Hot Pepper (Capsicum Annum L.) Landraces. Journal of Agricultural Sciences. 61(1): 19–36.

17. Bekele B, Petros P, Oljira T, Andargie M. Genotypes performance and genetic variability studies in Hot Pepper (Capsicum annum L.). Trends in current biology; 2023. DOI: 10.14719/tcb.2863

18. Mishra B.B., H.G. Kidan, K. Kibret, M. Assen and B. Eshetu. Soil and land resource inventory at Alemaya University research farm with reference to land evaluation for sustainable agricultural management and production: Synthesis of working papers, Soil Science Bulletin. Alemaya University, Ethiopia.

19. Simret Burga. Influence of inorganic nitrogen and potassium fertilizers on seed tuber yield and size distribution of potato (Solanum tuberosum L.). An. M. Sc Thesis Presented to the School of Graduate Studies of Haramaya University, Ethiopia. 65p.

20. EARO (Ethiopian Agricultural research Organization). Released crop varieties and their recommended cultural practices. Progress report. Addis Ababa, Ethiopia.

21. IPGRI. Descriptors for Capsicum (Capsicum sp.). Rome, Italy.

22. Gomez, K. A and Gomez, A. A. Statistical procedure for an agricultural research, 2nd edition. John Wily and Sons, USA. Pp: 680.

23. SAS Institute. SAS software. SAS Institute INC., Cary. NC. USA.

24. Cochran, W. G., and Cox, G. M. Experimental Designs. pp. 127–131. cooperation, Ministry of agriculture, Government of West Bengal. (online) http://dse.wb.nic.in/Home.htm.

25. Falconer DS, Mackay TFC. An Introduction to quantitative genetics. Ed, 4.Printice; 1996

26. Burton, G.W. Quantitative inheritance in grasses. Proceedings of 6th International Grassland Congress; 1952. 1: 227–283

27. Burton GW, De Vane EH. Estimating heritability in tall fescue (Fistvea arundiancea) from the replicated clonal material. Agriculture Journal; 1953. 1953;45: 284–291.

28. Sivasubramaniah, S and Menon, M. Heterosis and inbreeding depression in rice. Madras Agricultural Journal; 1973. 60: 11–39.

29. Hanson, C. H., Robinson, H. F. and Comstock, R. E. Biometrical studies of yield in segregating population of Korean lespedse. Agronomy Journal; 1956. 48, 267–282.

30. Lush, J.L. Inter-size correlation regression of offspring on dairy as a method of estimating heritability of characters. In: Proceedings American Society of Animal Production; 1940. 32: 293–301.

31. Robinson, H.F. Quantitative genetics in relation to breeding of the centennial of mendalism. Indian Journal of Genetics; 1966. 26, 171–187.

32. Johnson, H.W., Robinson, H.F. and Comstock, R.E. Genotypic and phenotypic correlations in soybeans and their implication in selection. Agronomy Journal; 1955. 47: 477–483.

33. Allard RW. Principles of Plant Breeding. John Willey and Sons; 1960, New York. p. 485.

34. Vijaya, H.M., Gowda, A.P. M. and Nehru, S. D. Genetic variability, correlation coefficient and path analysis in chilli (Capsicum annuum l.) genotypes. Environmental Life Science; 2014, 7 (3): 175–178.

35. Verma, S. K., Singh, R. K. and Arya, R. R. Genetic variability and correlation studies in chillies. Progressive Horticulture; 2004, 36(1): 113–117.

36. Bhagyalakshmi, P. V., Shankar, C. R., Subbramanyam, D. and Babu, V. G. Heterosis and combining ability studies in chillies. Indian Journal of Genetics and Plant Breeding; 1990, 51 (4): 420–423.

37. Ibrahim, M., Ganiger, V.M., and Yenjerappa, S.T. Genetic variability, heritability, genetic advance and correlation studies in chilli. Karnataka J Agril Sci; 2001, 14:784–787.

38. Janaki, M., Naram, L., Naidu, C., Ramana, V., and Rao, M. P. Assessment of Genetic Variability, Heritability and Genetic Advance for Quantitative Traits in Chilli (Capsicum annuum L.). International Quarterly Journal of Life Science; 2015, 10(2): 729–733.

39. Manju, P. R. and Sreelathakumary, I. Genetic Variability, Heritability and Genetic Advance in Hot Chilli (Capsicum Chinense Jacq.). Journal of Tropical Agriculture; 2002, 40: 4–6.

40. Khurana, D. S., Singh, P. and Hundal, J. S. Studies on genetic diversity for growth, yield and quality traits in chilli (Capsicum annum L.). Indian Journal of Horticulture; 2003, 60 (3): 277–282.

41. Singh, Y., Sharma, M. and Sharma, A. Genetic Variation, Association of Characters, and their Direct and Indirect Contributions for Improvement in Chilli Peppers, International Journal of Vegetable Science; 2009, 15(4): 340–368.

42. Sreelathakumary, I. and Rajamony, L. Variability, heritability and genetic advance in chilli (Capsicum annuum L.). Journal of Tropical Agriculture; 2015, 42(1-2): 35–37.

43. Maurya, A. K., Kushwaha, M. L., and Singh, B. K. Genetic Studies in Chilli (Capsicum annuum L.). International Journal of Science & Technology; 2015, 2321 – 919.

44. Comstock, R. E., and H. F. Robinson. The components of genetic variance in populations of biparental progenies and their use in estimating the average degree of dominance. Biometrics (1952): 254–266.

45. Sood, S., Sood, R., Sagar, V. & Sharma, K. C. Genetic Variation and Association Analysis for Fruit Yield, Agronomic and Quality Characters in Bell Pepper. International. Journal of Vegetable Science; 2009, 15 (3): 272–284.

